## Supplementary figures and images for "Horizontally transferred cell-free chromatin particles function as autonomous satellite genomes and vehicles for transposable elements within host cells"

### Supplementary Fig. S1

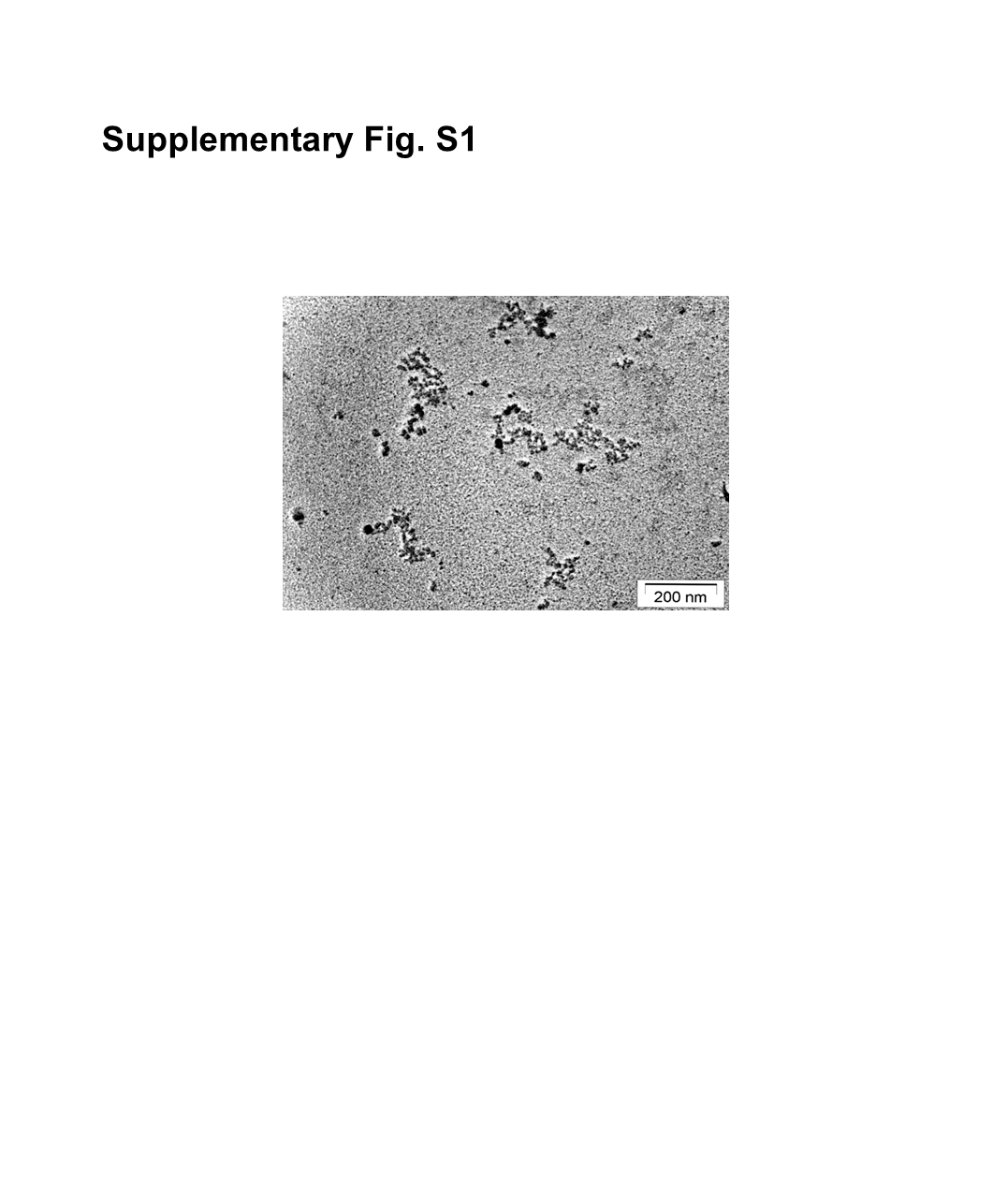

### Supplementary Fig. S2a

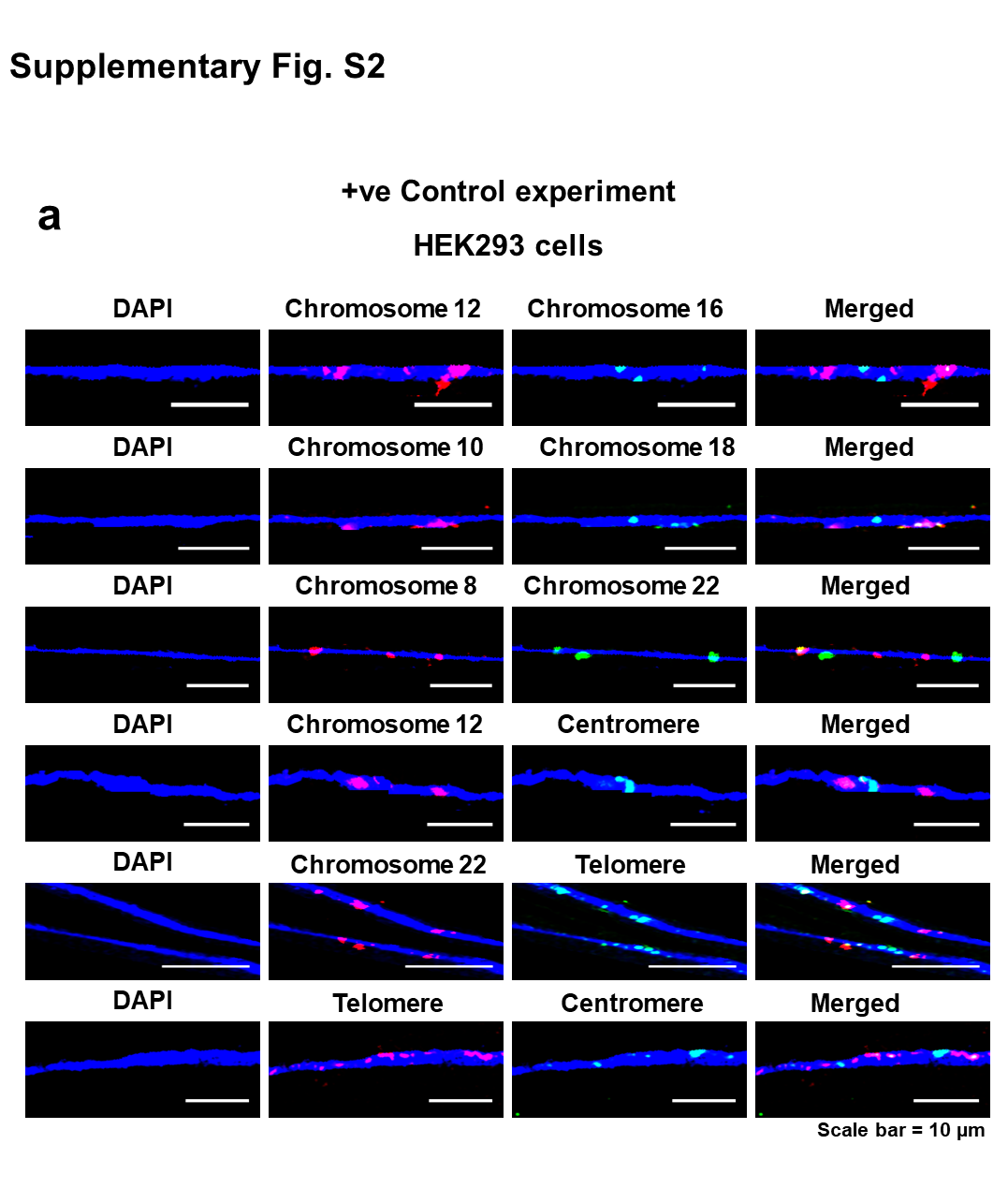

### Supplementary Fig. S2b

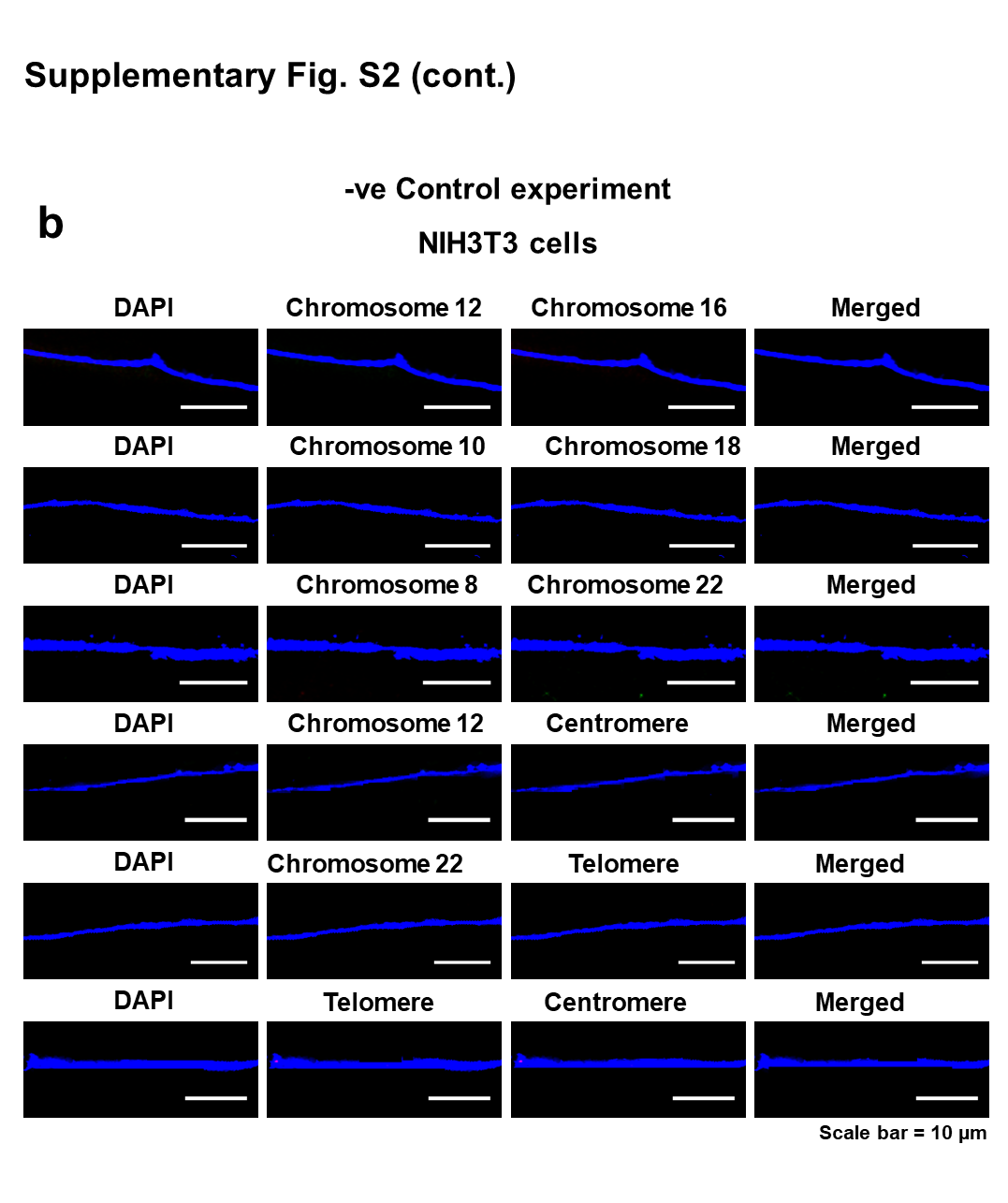

### Supplementary Fig. S3a-b

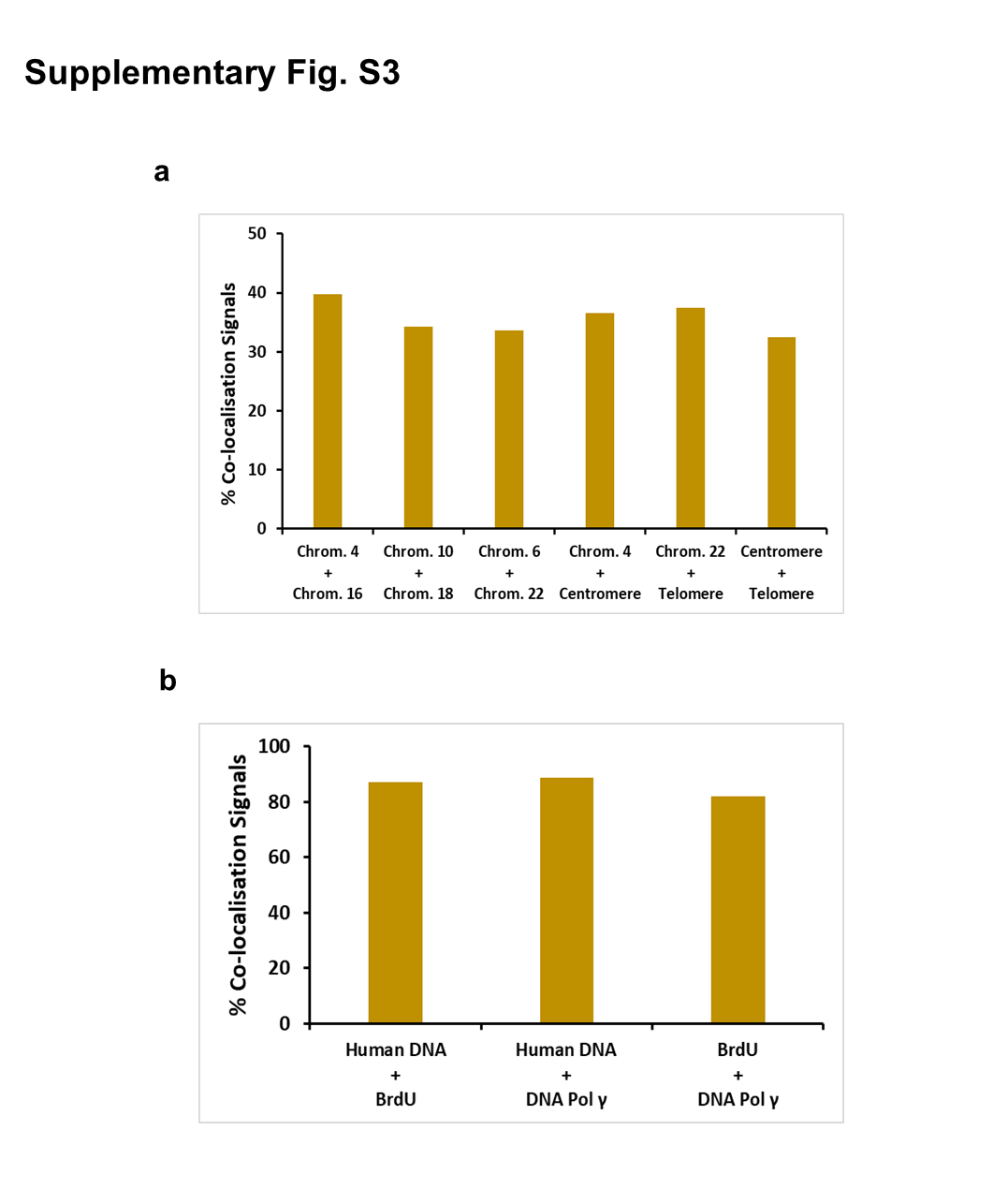

### Supplementary Fig. S3c-d

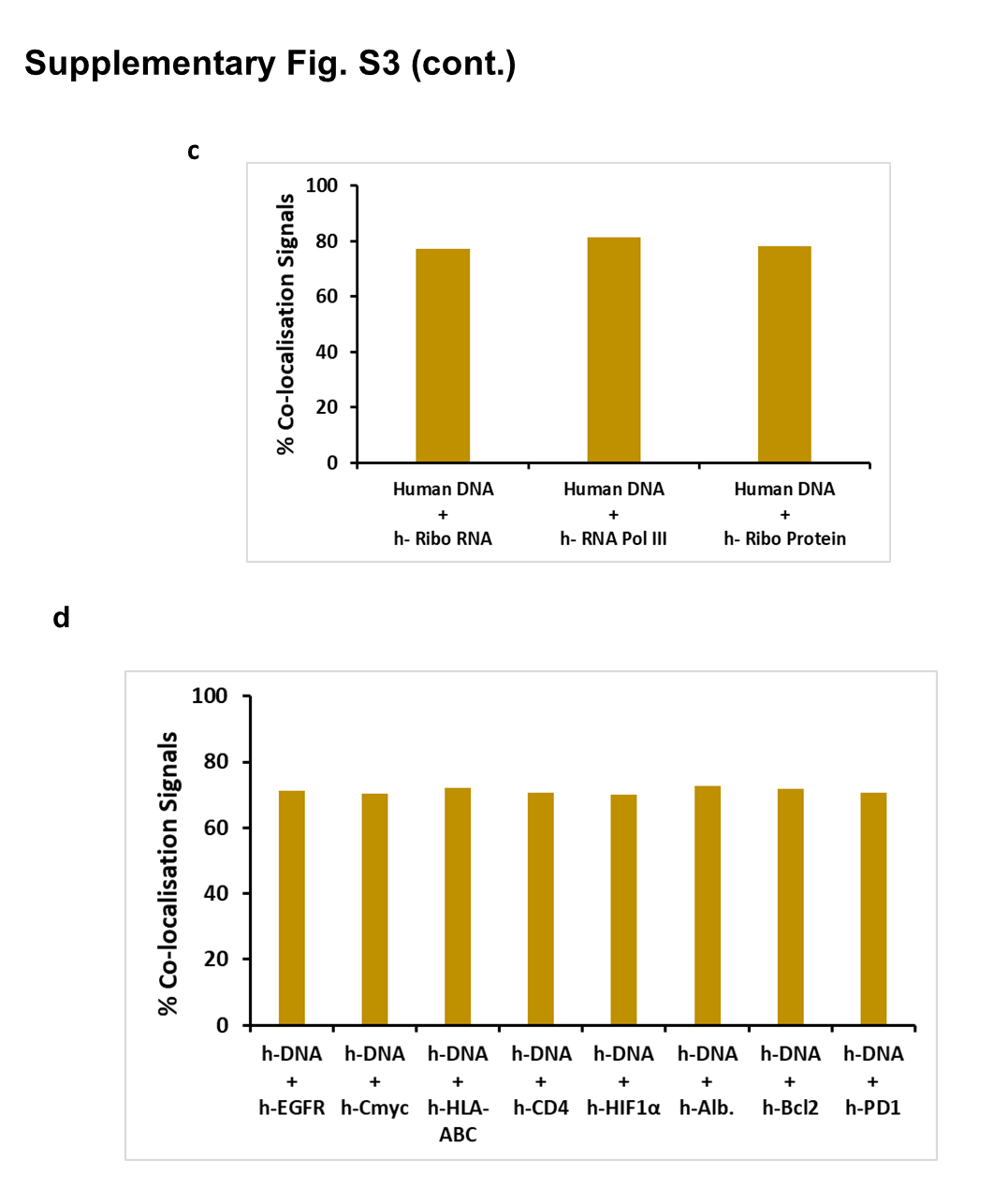

### Supplementary Fig. S3e-f

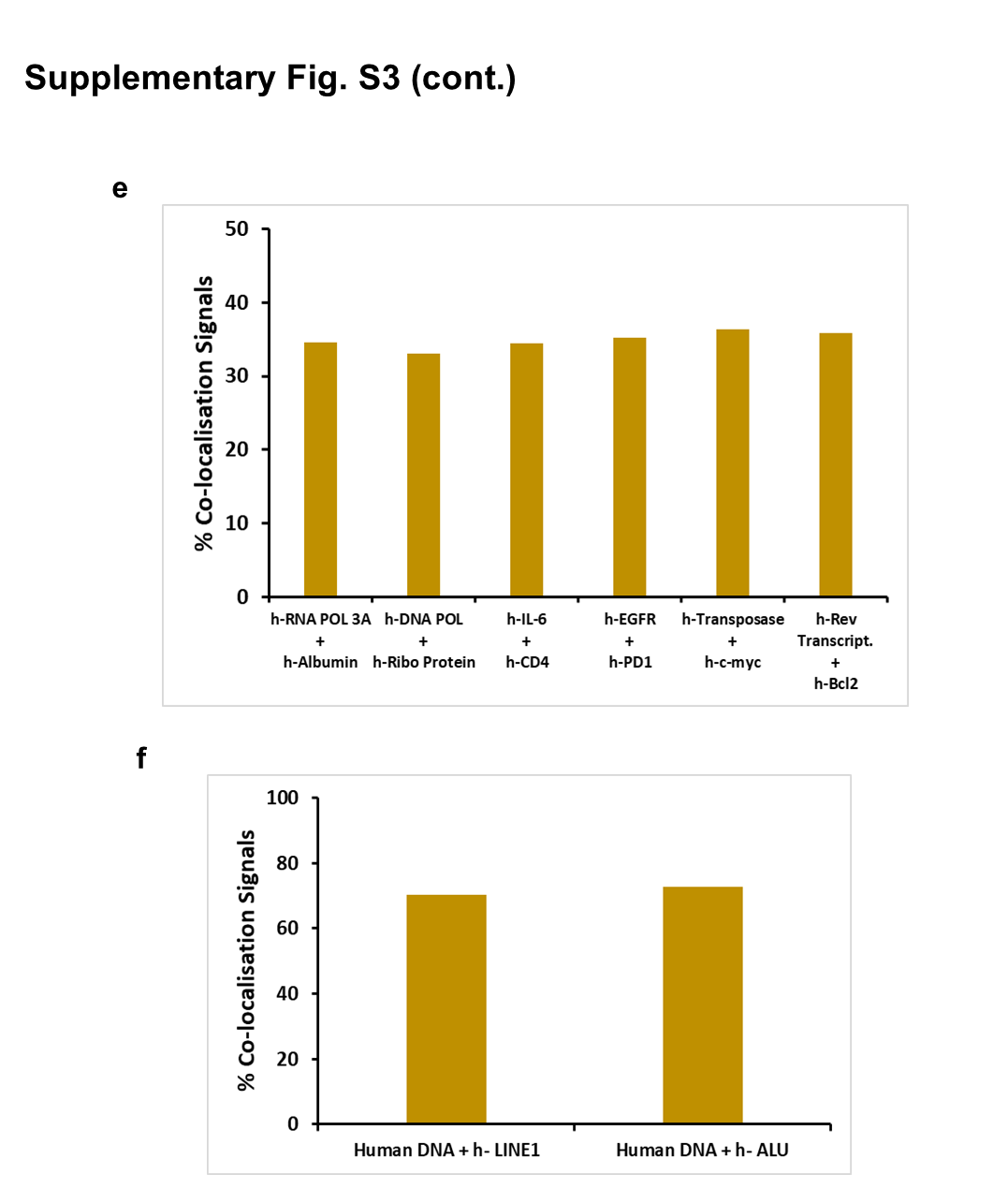

### Supplementary Fig. S3g-h

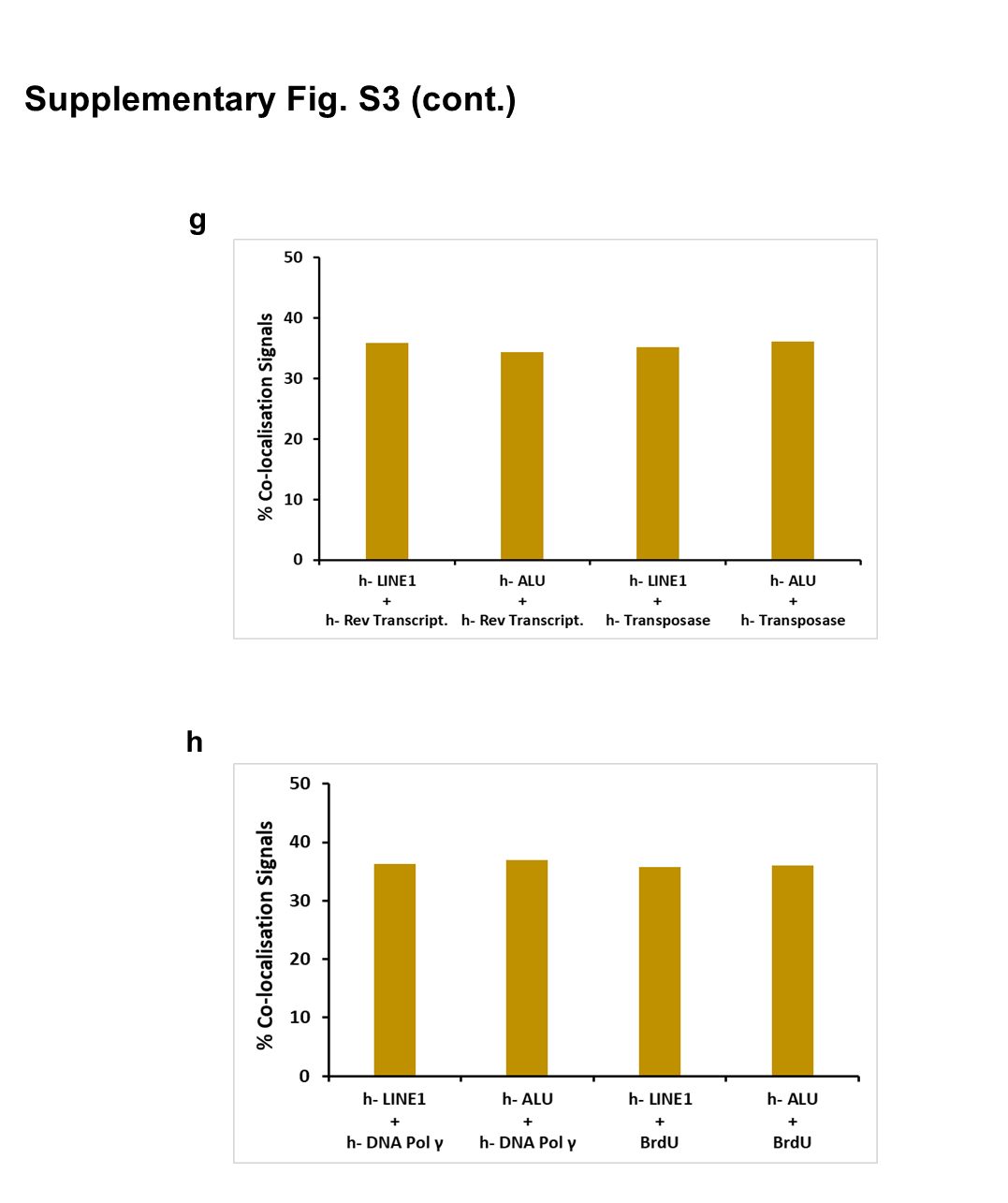

### Supplementary Fig. S4a-c

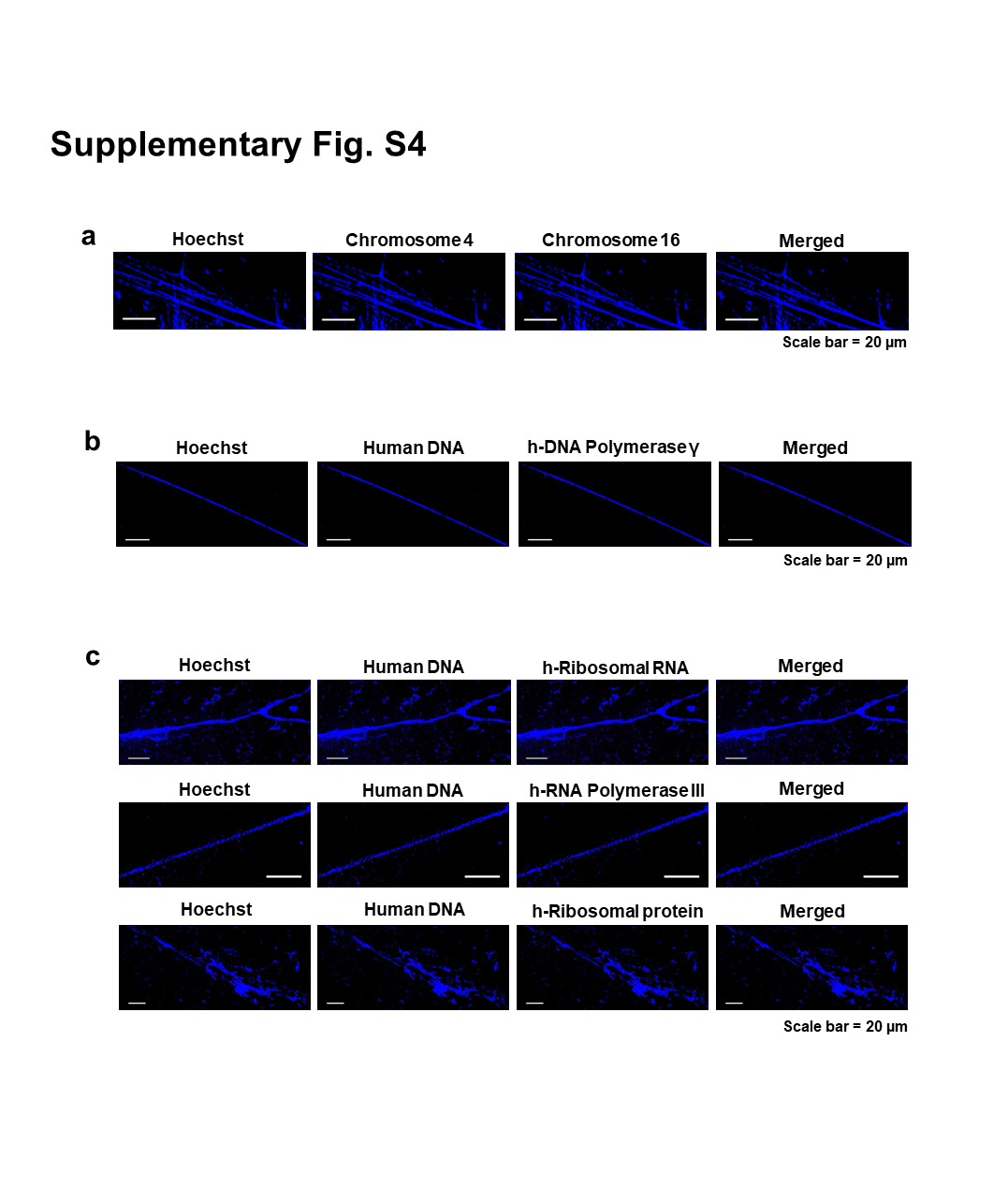

### Supplementary Fig. S4d

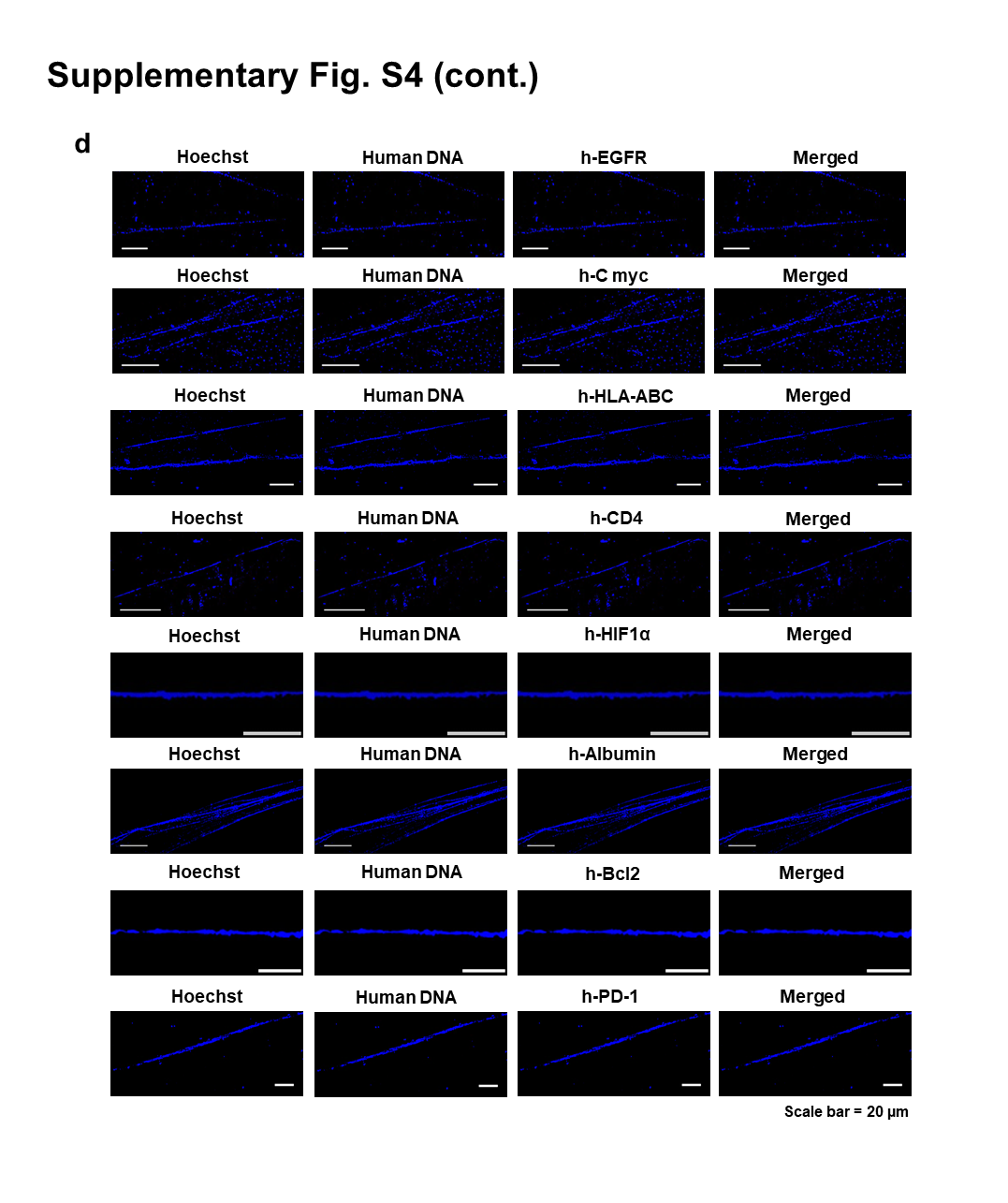

### Supplementary Fig. S4e-f

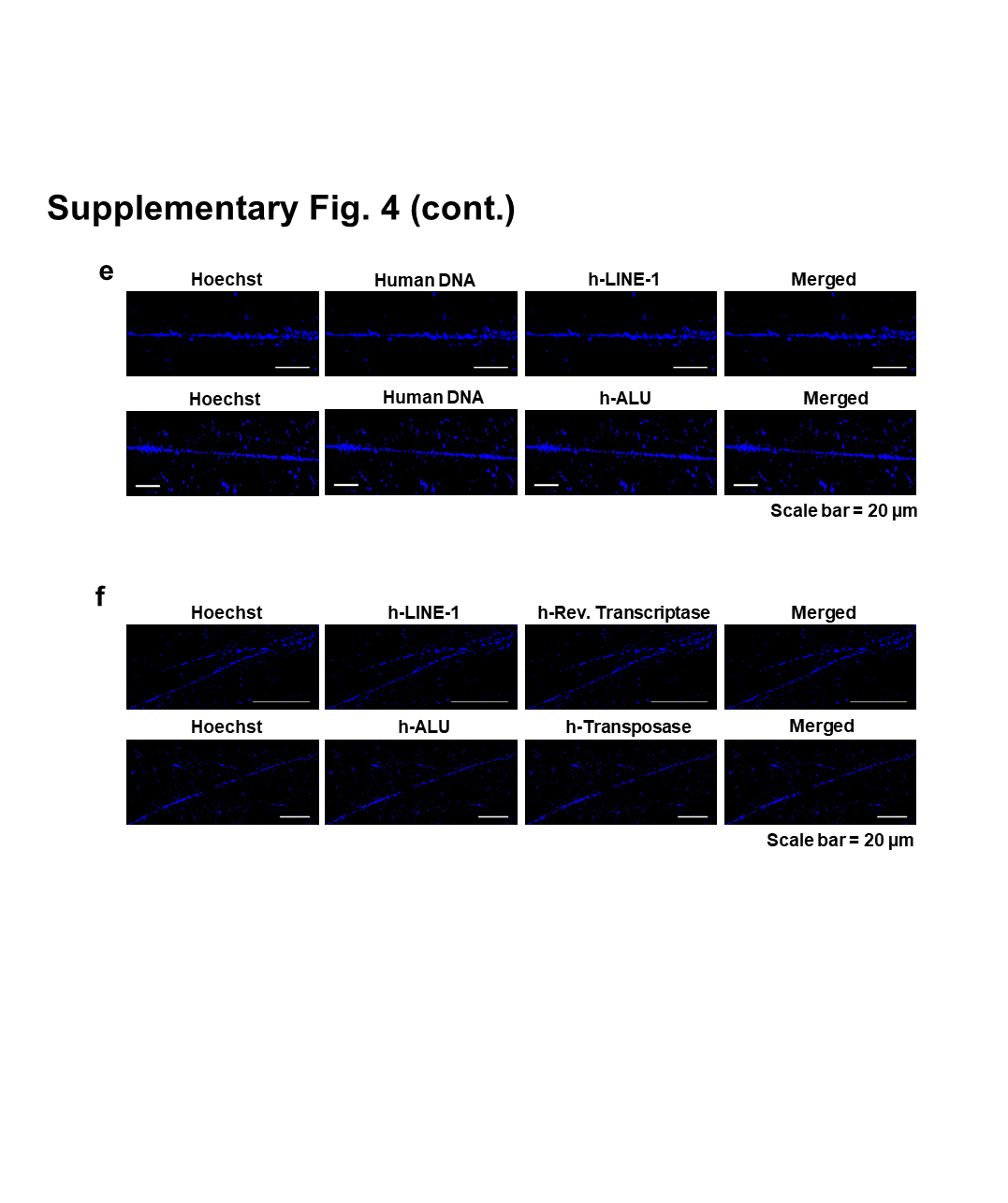

### Supplementary Fig. S5

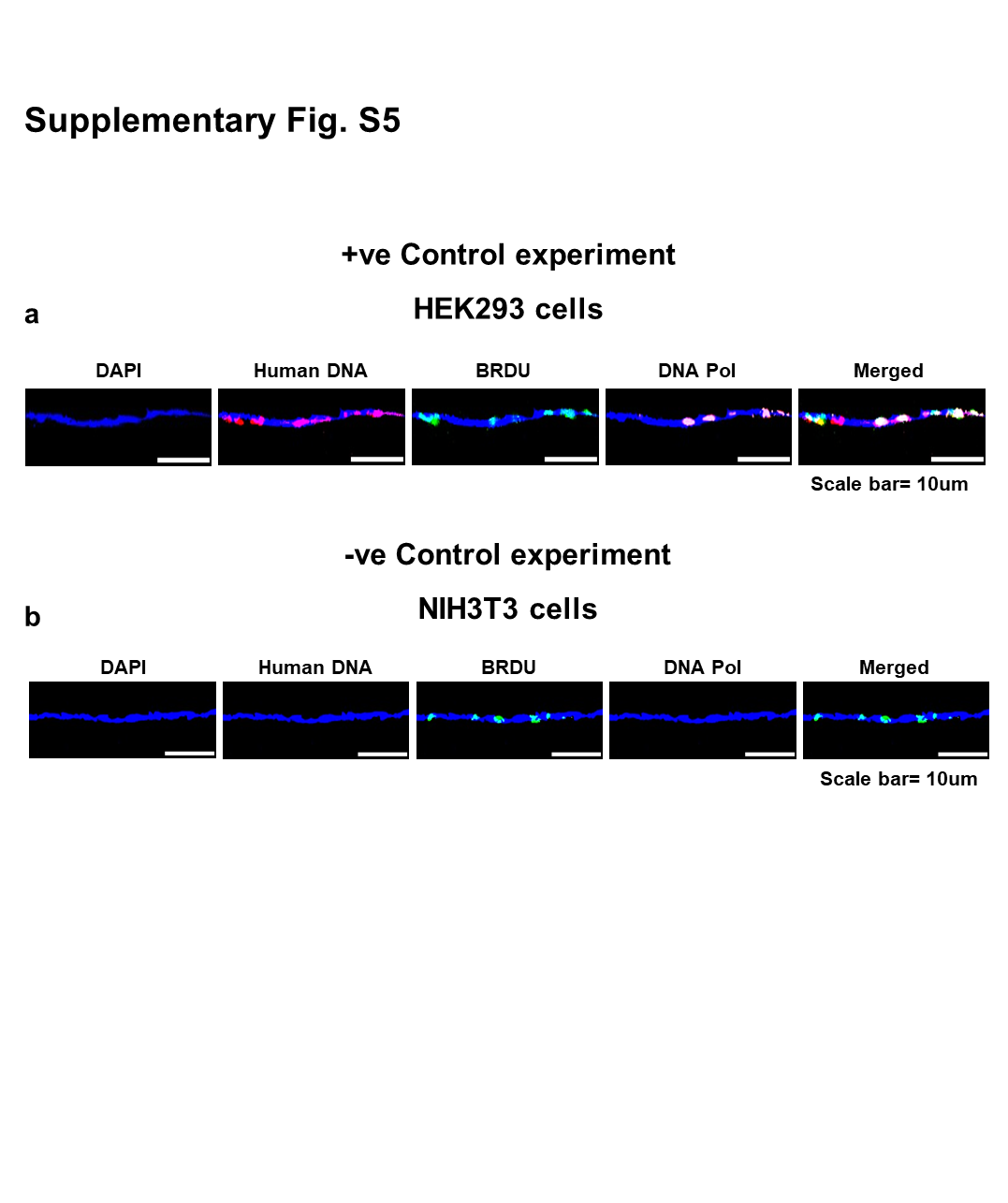

### Supplementary Fig. S6a-b

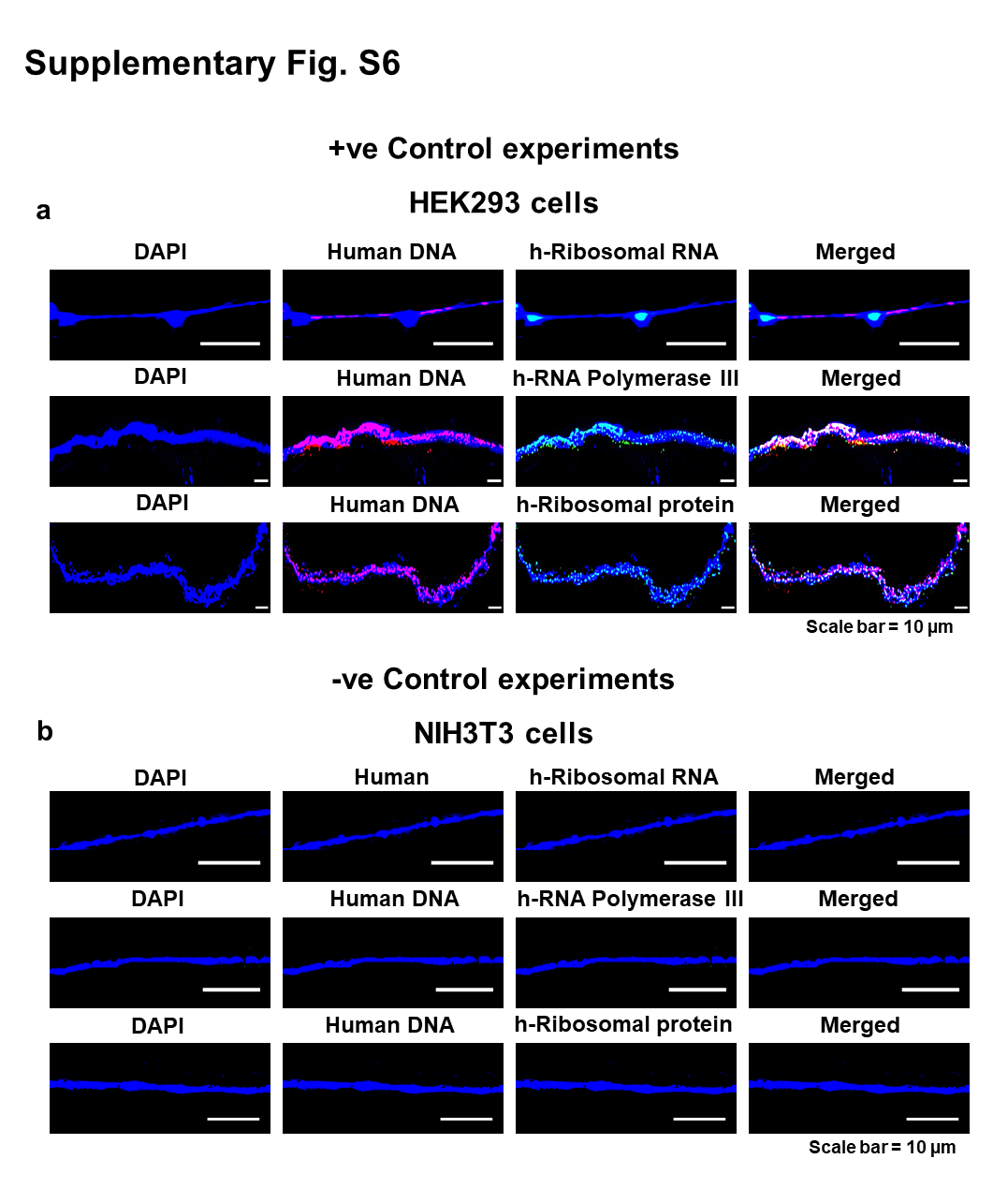

### Supplementary Fig. S7

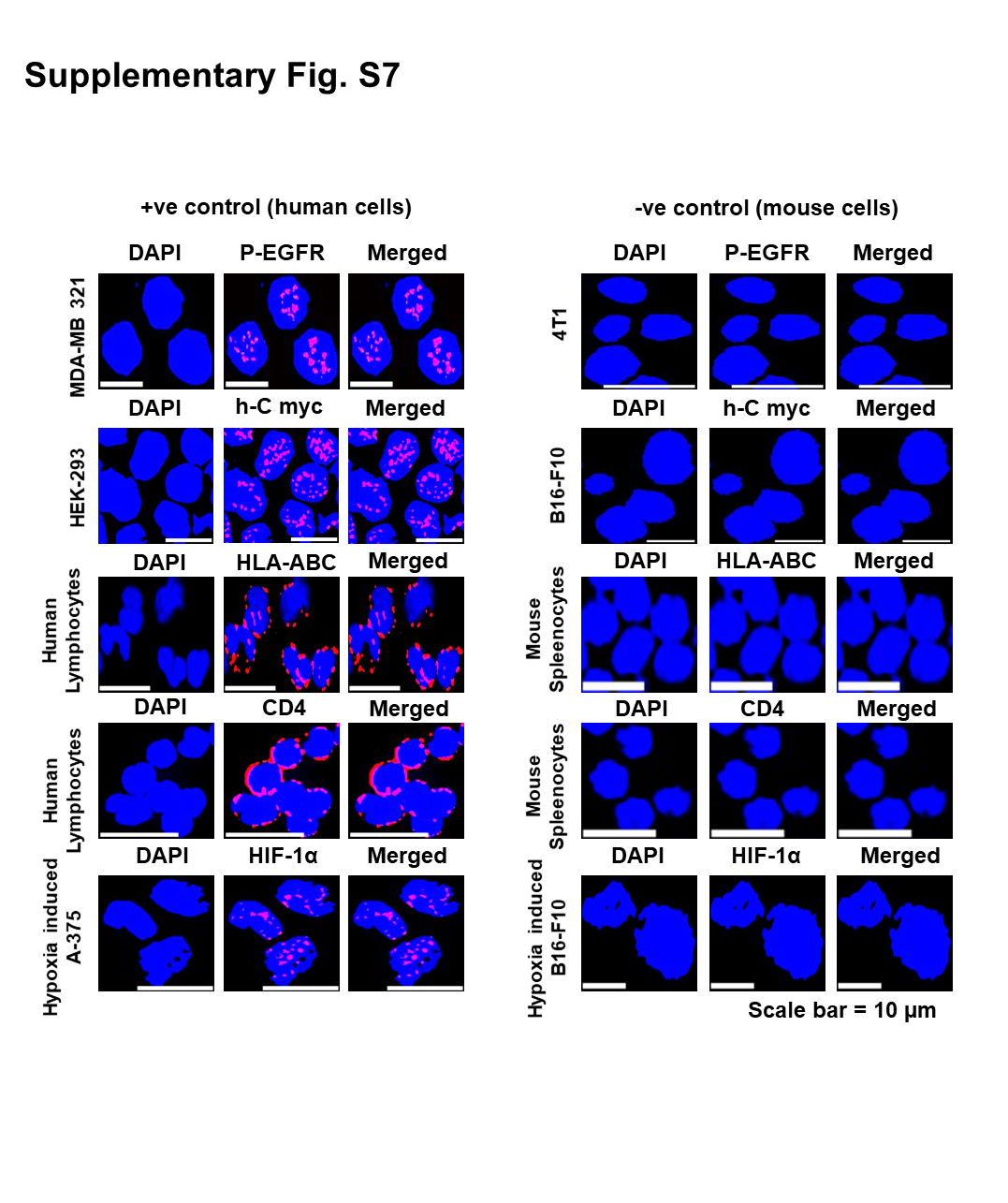

### Supplementary Fig. S7 (Cont..)

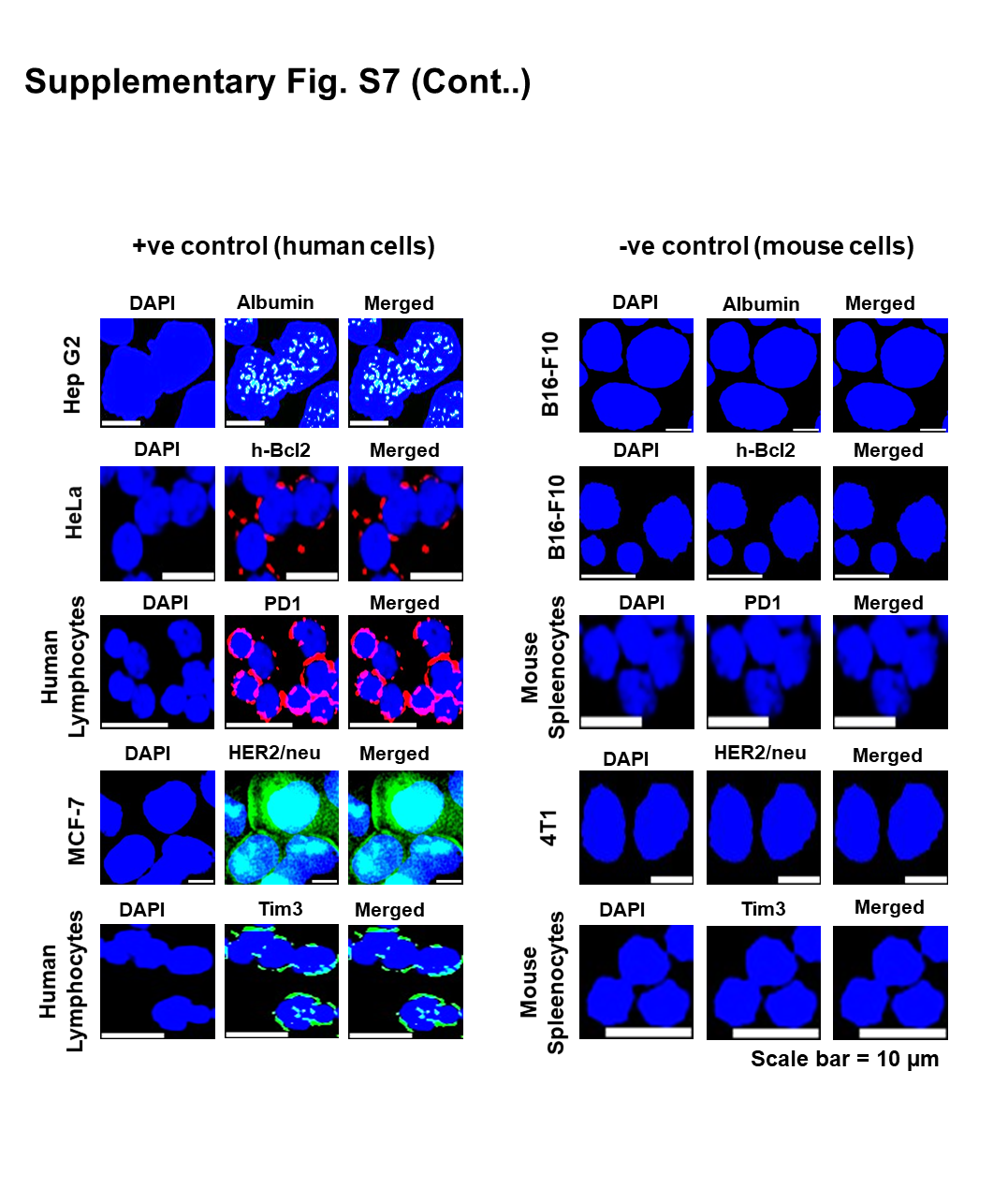

### Supplementary Fig. S8

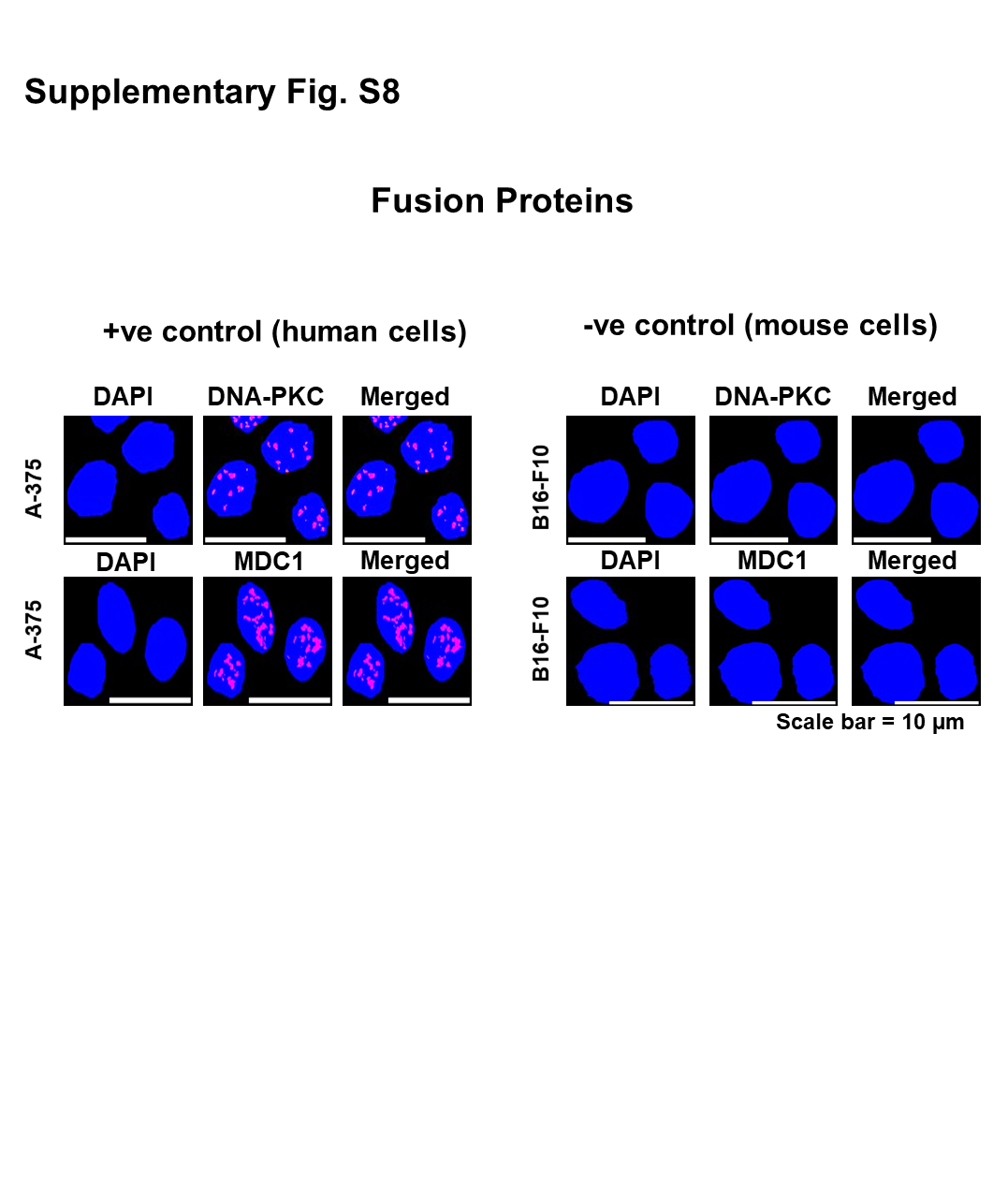

### Supplementary Fig. S9a

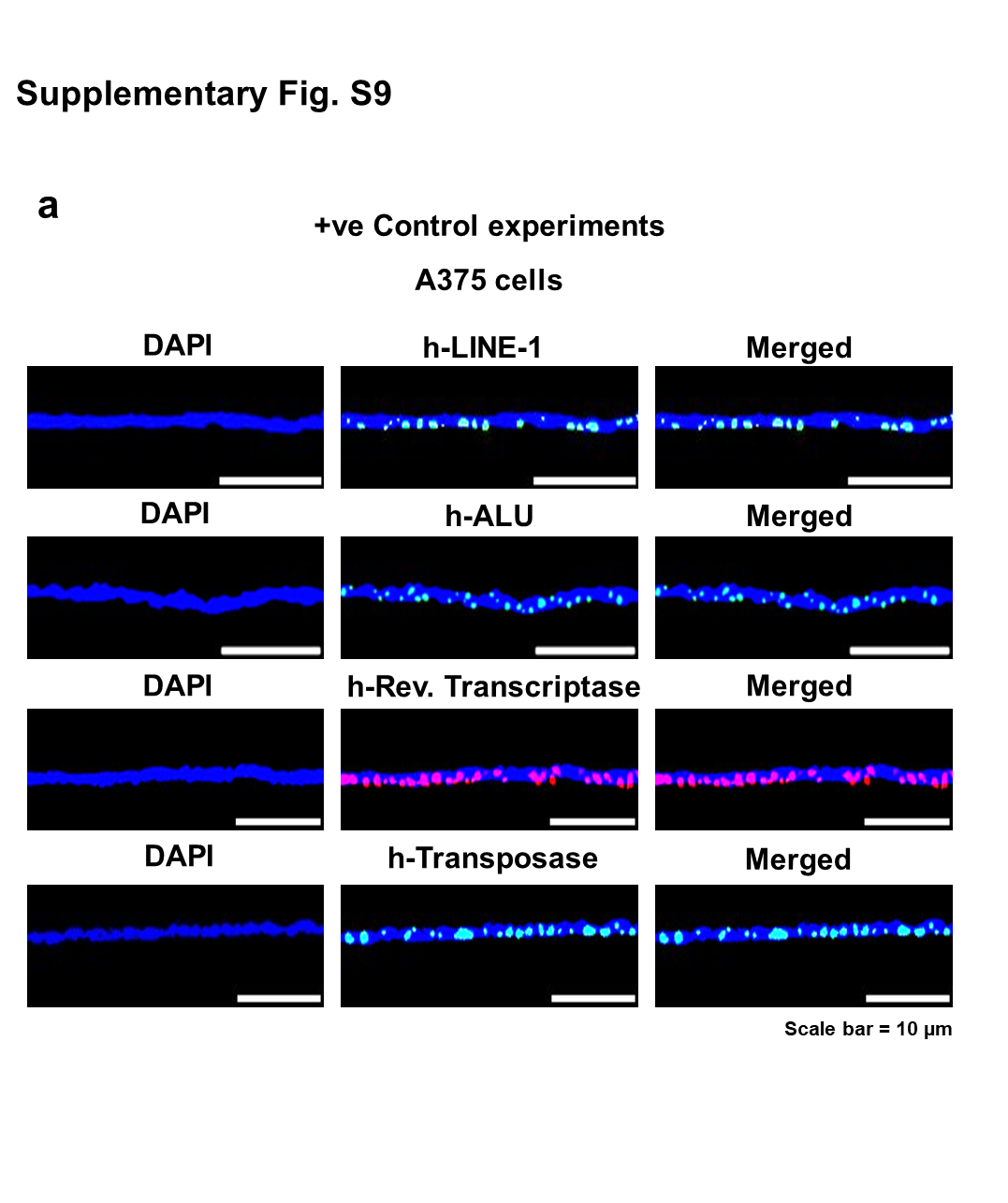

### Supplementary Fig. S9b

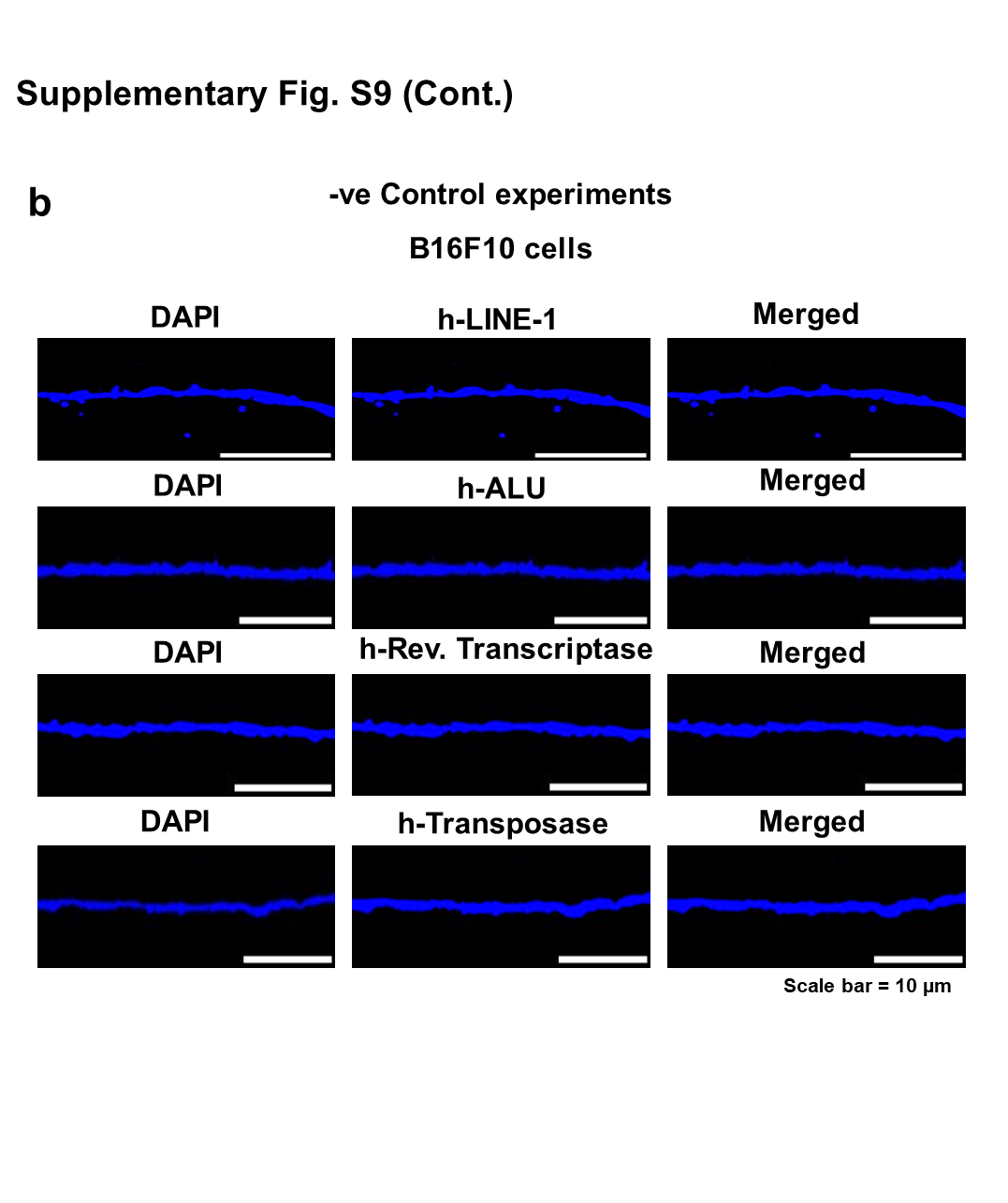

### Supplementary Fig. S10

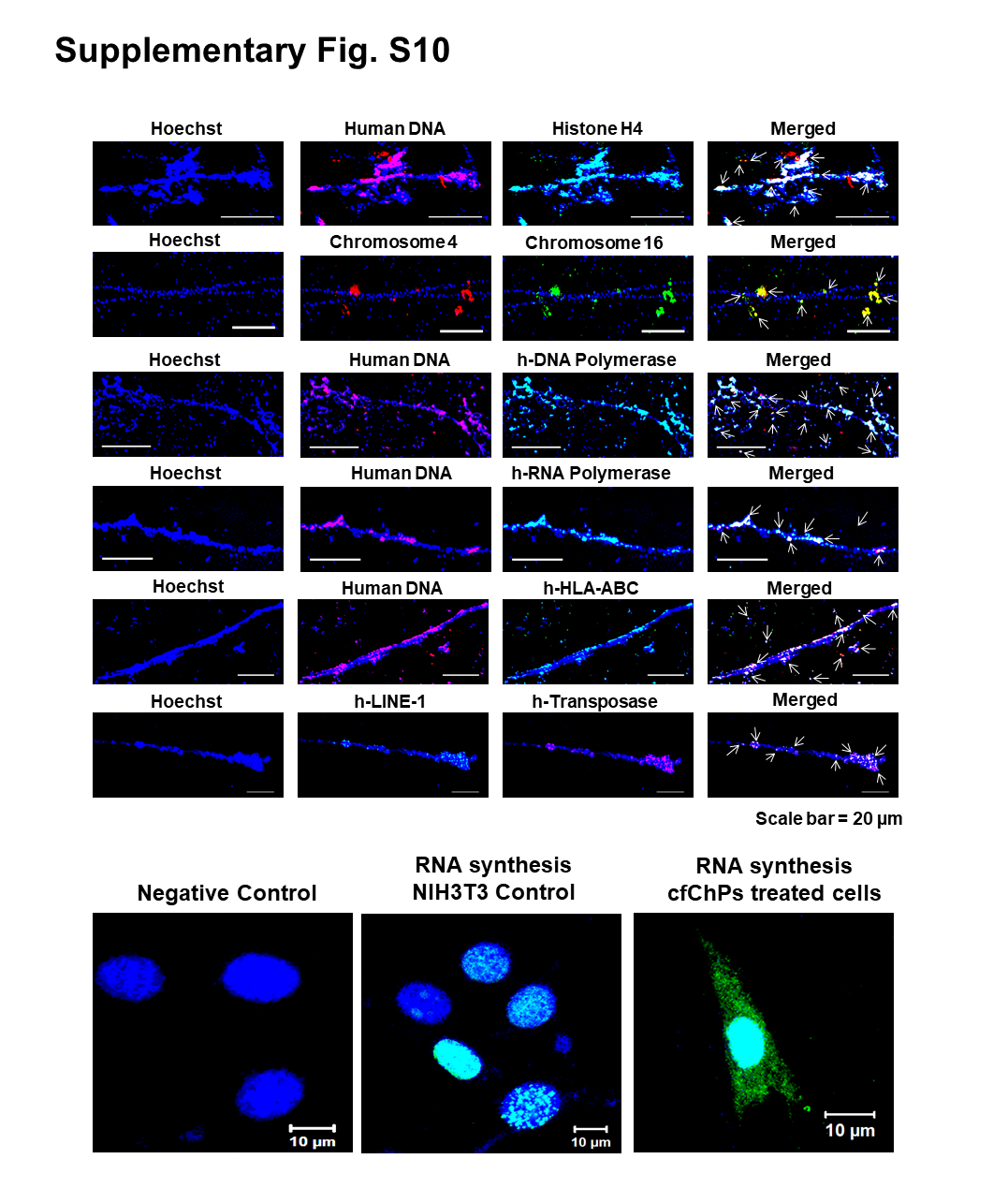

### Supplementary Fig. S11

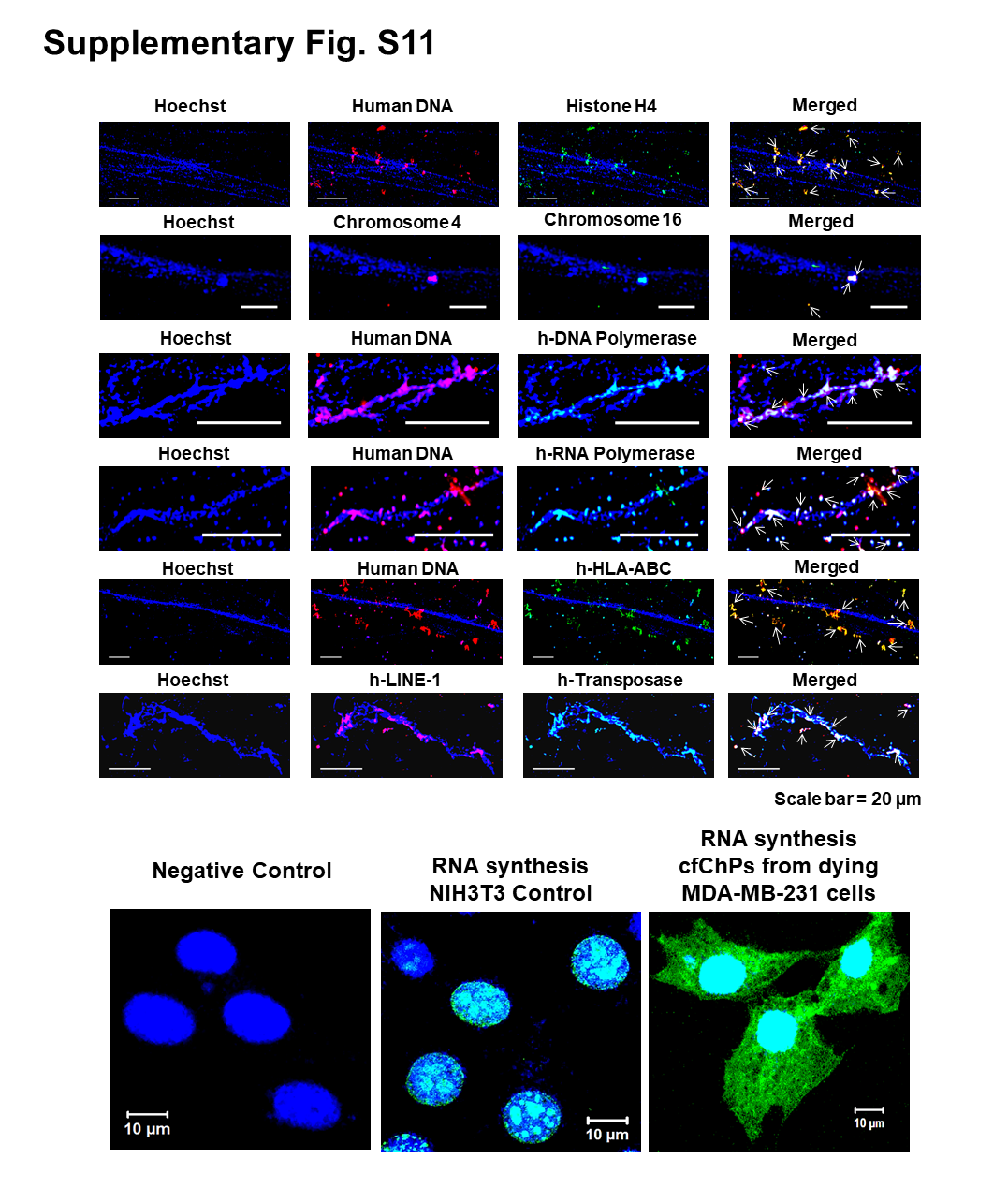

### Supplementary Fig. S12

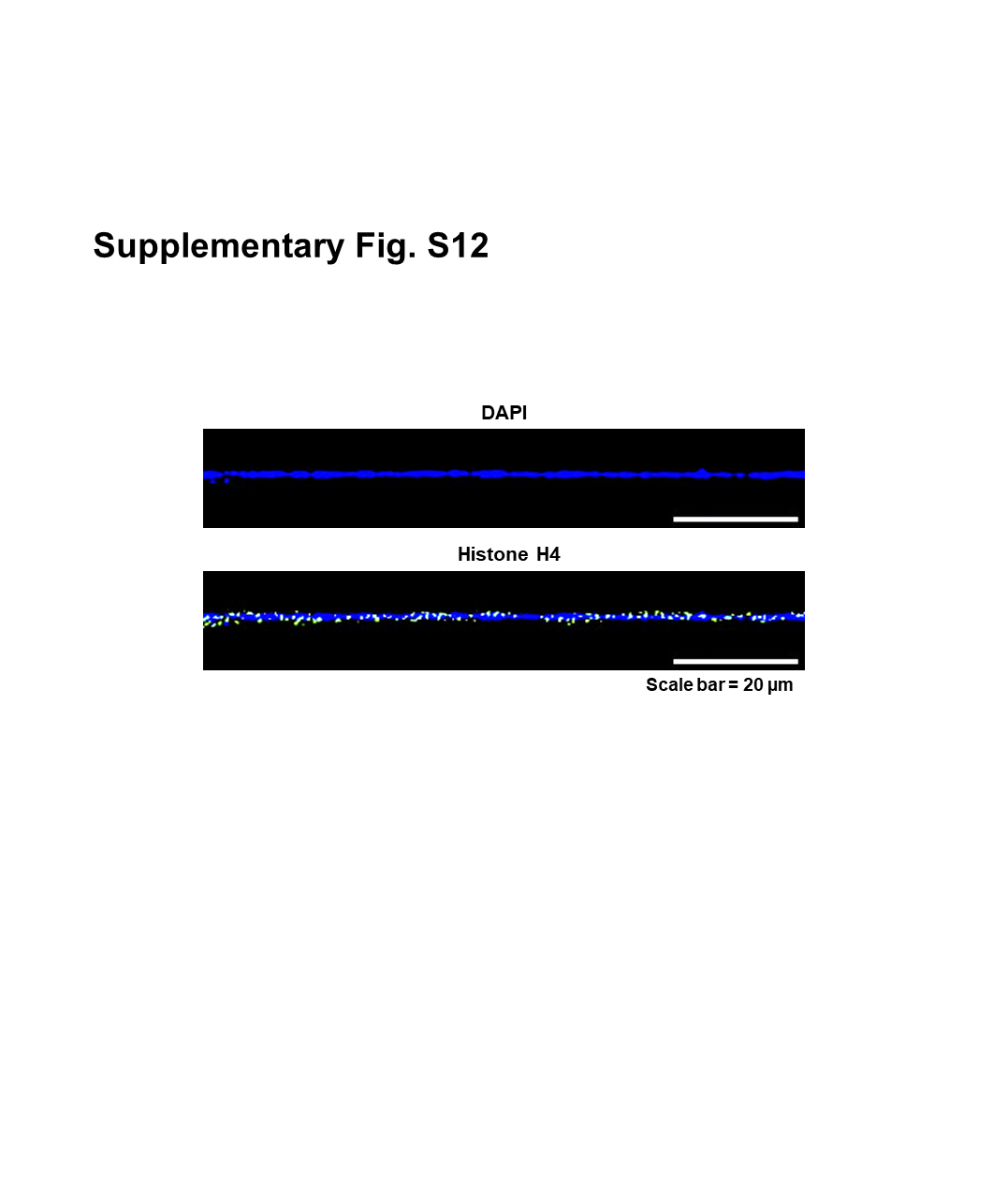

### Supplementary Fig. S13

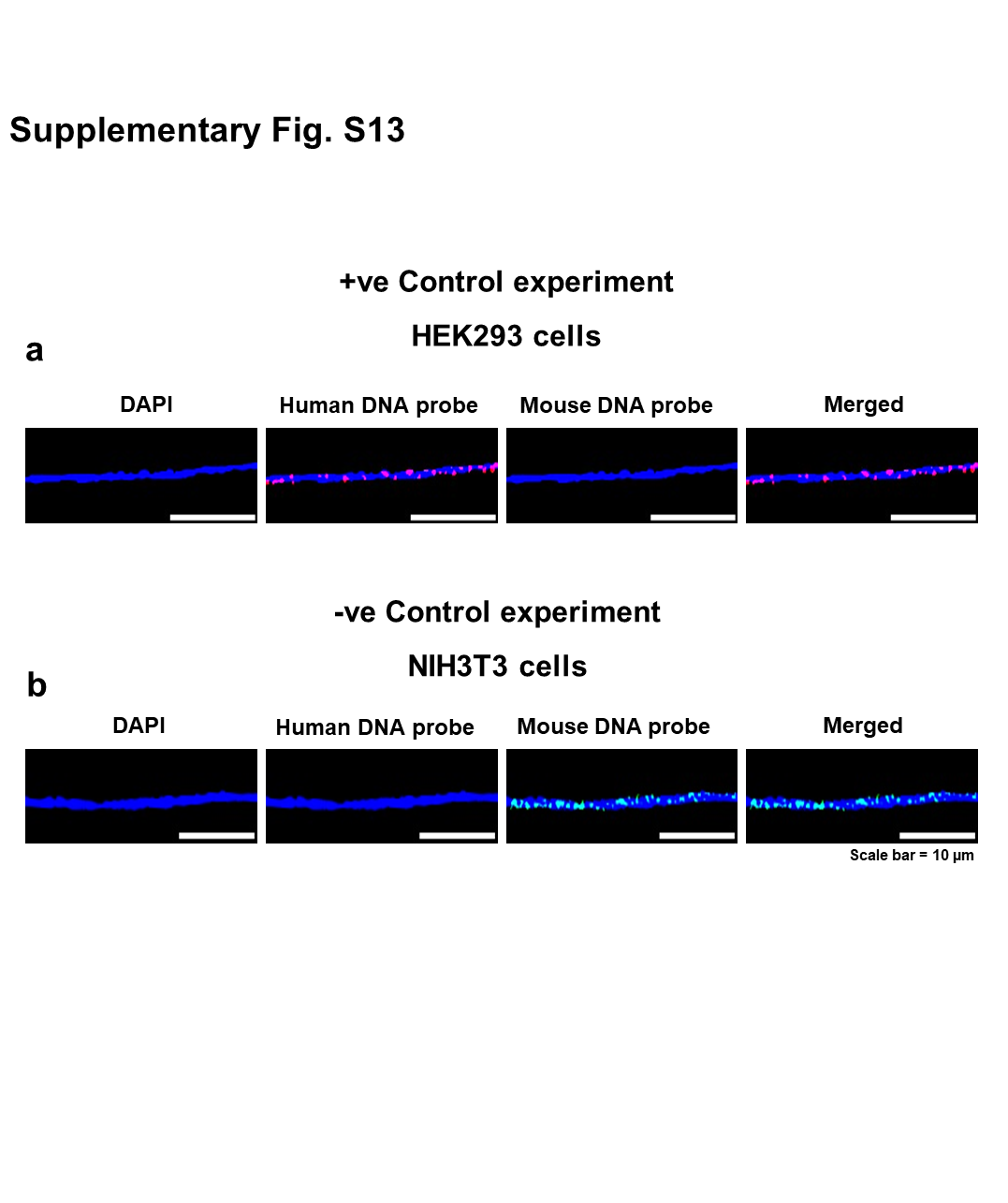

### Supplementary Fig. S14

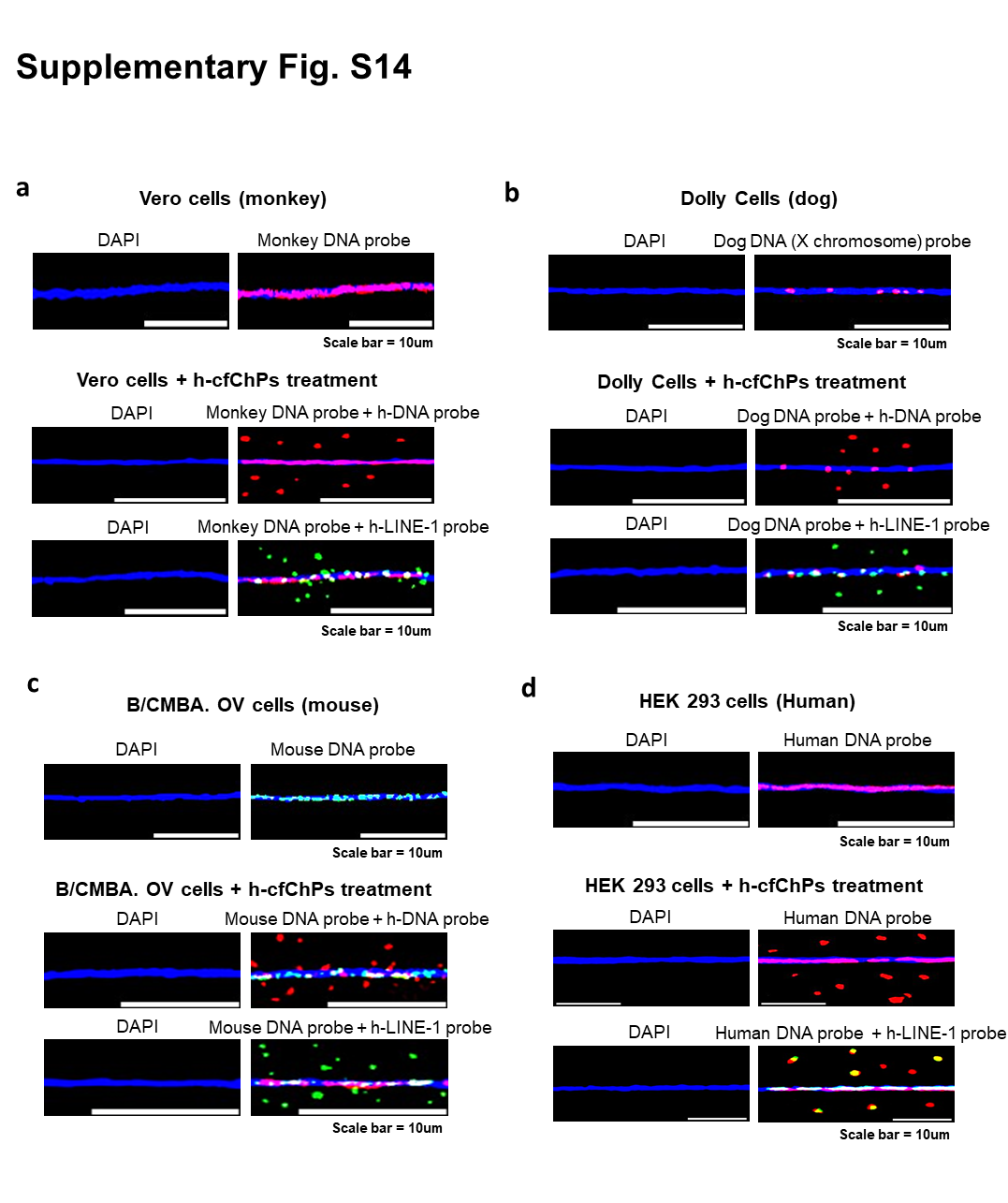

### Supplementary Fig. S15

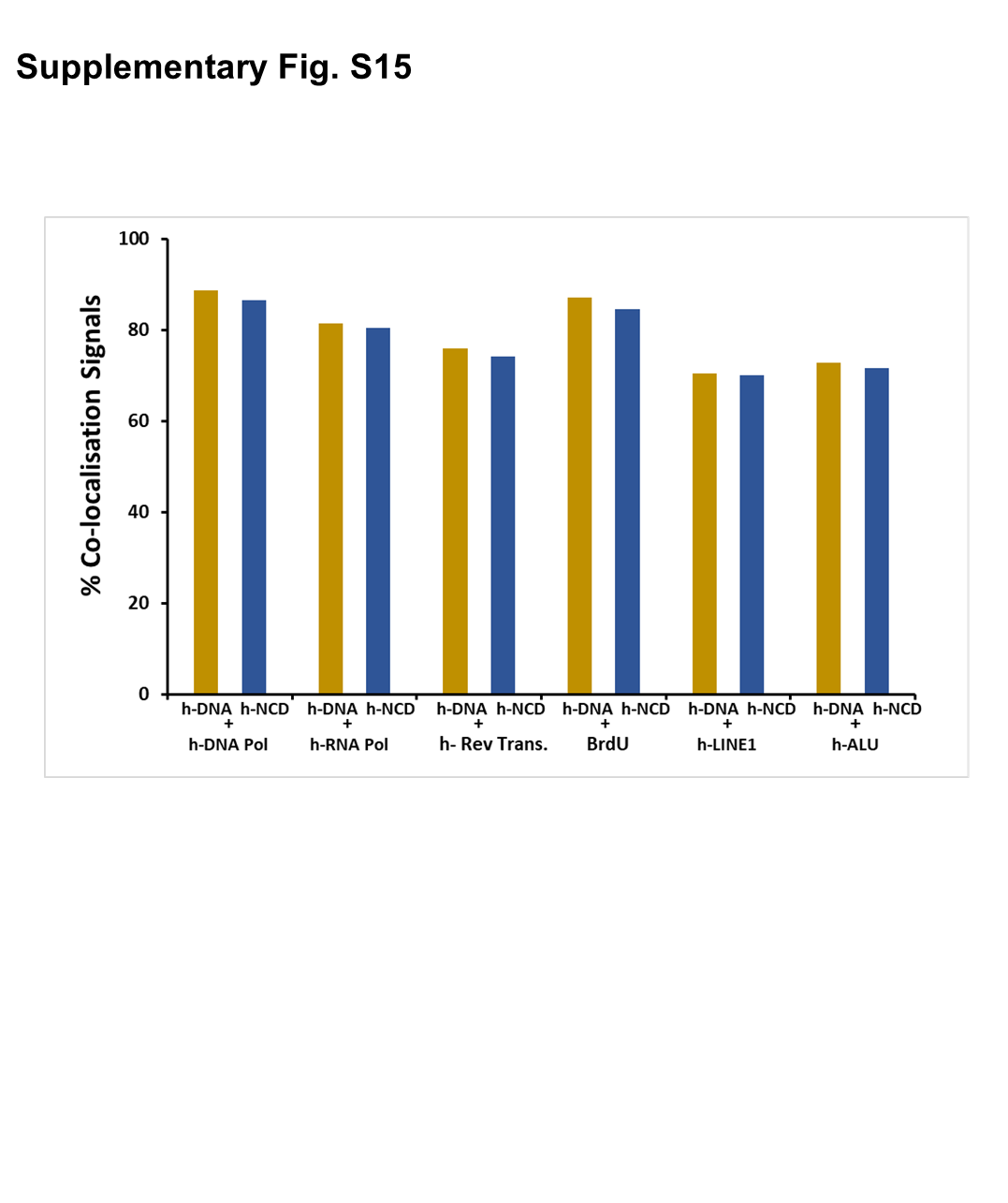
