## Supplementary material for "Horizontally transferred cell-free chromatin particles function as autonomous satellite genomes and vehicles for transposable elements within host cells": Legends to Supplementary Figures

**Legend to Supplementary Figures**

**Supplementary Figure S1.** Electron microscopy image of cfChPs isolated from pooled serum of patients with cancer. A ‘beads-on-a-string’ appearance typical of chromatin is clearly seen. Reproduced with permission from Mittra et al., 2015.

**Supplementary Figure S2.** Chromosome specific probes that were used are human specific. Chromatin fibres prepared from cfChPs treated passaged cells were probed with different pairs of chromosome specific FISH probes and those against telomere and centromere. **a.** Positive control cells using HEK-293 (human embryonic kidney) cells; **b.** Negative control experiments using NIH3T3 (mouse embryonic fibroblast) cells.

**Supplementary Figure S3.** Histograms representing quantitative results of the degree of co-localisation of fluorescent signals (red and green) of different pairs of biomarkers. Five hundred fluorescent signals were analysed and the number of co-localising red and green signals was estimated. The results were expressed in percentage terms **(a-h)**.

**Supplementary Fig. S4.** Control experiments to detect human DNA and/or proteins in control NIH3T3 cells that had not been exposed to cfChPs. Neither the human specific FISH probes nor the human specific antibodies detected any human DNA or protein signals in mouse cells **(a – f)**.

**Supplementary Fig. S5.** Control experiments to show that the antibody used against DNA Polymerase γ was human specific and did not cross-react with mouse. **a.** Positive control cells using HEK-293 (human embryonic kidney) cells; **b.** Negative control experiments using NIH3T3 (mouse embryonic fibroblast) cells.

**Supplementary Fig. S6.** Control experiments to show that the FISH probes and antibodies used against the components of the protein synthetic machinery viz. ribosomal RNA, RNA polymerase and ribosomal protein were human specific and did not cross-react with mouse. **a.** Positive control cells using HEK-293 (human embryonic kidney) cells; **b.** Negative control experiments using NIH3T3 (mouse embryonic fibroblast) cells.

**Supplementary Figure S7.** Control experiments to show that the antibodies used against the various proteins were human specific and did not cross-react with mouse. MDA-MB-231 (human breast cancer cells); HEK-293 (human embryonic kidney); A-375 (human melanoma cells); 4T1 (mouse mammary cancer cells); B16F10 (mouse melanoma cells). A-375 and B16F10 cells were kept in hypoxic chamber (1% O_2_) for 48h to induce activation of HIF1-α. Hep G2 (human hepatocellular carcinoma cells); HeLa (human cervical camcer cells); B16F10 (mouse melanoma cells); 4T1 (mouse mammary cancer cells).

**Supplementary Figure S8.** Control experiments to show that the antibodies used against the components of fusion proteins that were not included in Supplementary Figure S7 were human specific and did not cross-react with mouse. A-375 (human melanoma cells); B16F10 (mouse melanoma cells).

**Supplementary Fig. S9.** Control experiments to show that the probes used against LINE-1 and *Alu* transposable elements and the antibodies against reverse transcriptase and transposase were human specific and did not cross-react with mouse. cfChPs treated cells in continuous passage were used to study species specificity of the probes and antibodies. **a.** Positive control cells using A-375 (human melanoma) cells; **b.** Negative control experiments using B16F10 (mouse melanoma) cells.

**Supplementary Figure S10.** NIH3T3 cells treated with cfChPs isolated from the serum of healthy individuals and which were kept in continuous passage show similar characteristics and properties as those of cfChPs isolated from cancer patients.

**Supplementary Fig. S11.** NIH3T3 cells treated with conditioned medium containing cfChPs released from dying MDA-MB-231 breast cancer cells and which were kept in continuous passage show similar characteristics and properties as those of cfChPs isolated from cancer patients and healthy individuals.

**Supplementary Figure S12.** Representative image of a chromatin fibre prepared from untreated NIH3T3 cells stained with DAPI and anti-histone H4 antibody.

**Supplementary Figure S13.** Control experiments to show that the genomic DNA FISH probe used was human specific and did not cross-react with mouse. **a.** Positive control cells using HEK-293 (human embryonic kidney) cells; **b.** Negative control experiments using NIH3T3 (mouse embryonic fibroblast) cells.

**Supplementary Figure S14.** NIH3T3 cells are not unique in their ability to internalise cfChPs. Four different cell lines other than NIH3T3 cells were treated with cfChPs derived from human serum and chromatin fibres were prepared at 5^th^ passage. The chromatin fibres were probed with a whole genomic DNA probe or with a human specific LINE-1 probe. Human DNA and LINE-1 signal are clearly seen in the treated cells.

**Supplementary Figure S15.** Biological activities of the concatemers are attributable to non-coding DNA. Histograms showing the degree of co-localisation of fluorescent signals of human non-coding DNA and human whole genomic DNA and those of various biomarkers and LINE-1 and Alu elements. The degree of co-localization with the biomarkers was similar irrespective of whether a whole human genomic probe or one against non-coding DNA probe was used. These data suggested that biological activities of the concatemers are attributable to non-coding DNA.
