## Supplementary Tables for "Horizontally transferred cell-free chromatin particles function as autonomous satellite genomes and vehicles for transposable elements within host cells"

**Supplementary Table S1**

Clinical and demographic information of cancer patients and healthy individuals who provided blood samples for isolation of cell-free chromatin particles.

**Cancer patients**

| **Age** | **Sex** | **Diagnosis** |
| --- | --- | --- |
| 25 years | Female | Breast cancer |
| 35 years | Male | Tongue cancer |
| 22 years | Male | B-ALL |
| 32 years | Male | CML |
| 38 years | Male | Buccal mucosa cancer |

B-ALL = B cell acute lymphocytic lymphoma; CML = chronic myeloid leukaemia

**Healthy individuals**

| **Age** | **Sex** |
| --- | --- |
| 34 years | Male |
| 29 years | Female |
| 22 years | Male |
| 22 years | Male |
| 27 years | Male |

**Supplementary Table S2.**

**Antibodies and FISH probes used in this study**

**(As per vendors’ specifications)**

**Primary antibodies:**

| **Antibody** | **Host** | **Reactivity** | **Catalogue No.** | **Company** |
| --- | --- | --- | --- | --- |
| Histone H4 | Rabbit | Mouse, human | Custom-synthesised | Bioklone Biotech Pvt Ltd., India |
| Anti-BrdU | Rat | Human, mouse | ab6326 | Abcam, UK |
| DNA polymerase-γ | Rabbit | Rat, human | GTX100398 | GeneTex, USA |
| H3K4me3 | Mouse | Mouse, rat, human | ab1012 | Abcam, UK |
| H3K9me3 | Rabbit | Mouse, rat, human | #13969S | Cell Signaling Technologies, USA |
| Anti-DNA antibody | Mouse | Human, mouse, rat | NB110-89473 | Novus Biologicals LLC, USA |
| RNA polymerase  III A | Rabbit | Human | GTX106241 | GeneTex, USA |
| Ribosomal protein (RPLP0) | Mouse | Human | SAB1402899 | Sigma-Aldrich, USA |
| pEGFR | Rabbit | Human | 4404S | Cell Signalling Technology, USA |
| c-Myc | Mouse | Human | M4439 | Merck-Millipore Sigma, Germany |
| HLA-ABC class I | Mouse | Human | ab70328 | Abcam, UK |
| CD4 | Rabbit | Human | ab133616 | Abcam, UK |
| HIF-1α | Mouse | Human | 610959 | BD Biosciences, US |
| ALB (serum albumin) | Mouse | Human | sc-51515 | Santa-Cruz Biotechnology, USA |
| Bcl-2 | Mouse | Human | ab692 | Abcam, UK |
| PD-1 | Mouse | Human | ab52587 | Abcam, UK |
| HER2/NEU2 | Rabbit | Human | A0485 | DAKO- Agilent, USA |
| TIM-3 | Goat | Human | ab47997 | Abcam, UK |
| IL-6 | Rabbit | Human, Dog | ab6672 | Abcam, UK |
| Reverse transcriptase | Rabbit | Human | ab111584 | Abcam, UK |
| MDC-1 | Rabbit | Human | ab11169 | Abcam, UK |
| DNA PKCs | Rabbit | Human | ab32566 | Abcam, UK |
| Transposase | Rabbit | Human | MABE-1987 | Merck-Millipore Sigma, Germany |

**Secondary antibodies:**

| **Antibody** | **Catalogue No.** | **Company** |
| --- | --- | --- |
| Rabbit Anti-Goat IgG H&L (FITC) | ab6737 | Abcam, UK |
| Goat anti-mouse IgG H& L (FITC) | ab6785 | Abcam, UK |
| Goat Anti-Rabbit IgG (H+L)  TRITC | ab6718 | Abcam, UK |
| Goat Anti-Rabbit IgG (H+L)  FITC | AP307F | Merck-Millipore Sigma, Germany |
| Goat anti-Mouse IgG (H+L) Secondary Antibody, DyLight 633 | 35512 | Thermo-Fisher Scientific, USA |
| Goat anti-Rabbit IgG (H+L) Secondary Antibody, DyLight 633 | 35562 | Thermo-Fisher Scientific, USA |

**FISH probes:**

| **Probes** | **Catalogue no.** | **Company** |
| --- | --- | --- |
| Human genomic DNA (Cy3) | Custom-synthesised | Applied Spectral Imaging, Israel |
| Mouse genomic DNA (FITC) | Custom-synthesised | Applied Spectral Imaging, Israel |
| Human single-chromosome paint probes,  Chromosome 4 Red (Cy3) | Custom-synthesised | Applied Spectral Imaging, Israel |
| Human single-chromosome paint probes,  Chromosome 6 Red (Cy3) | Custom-synthesised | Applied Spectral Imaging, Israel |
| Human single-chromosome paint probes,  Chromosome 10 Red (Cy3) | Custom-synthesised | Applied Spectral Imaging, Israel |
| Human single-chromosome paint probes,  Chromosome 16 Green (FITC) | Custom-synthesised | Applied Spectral Imaging, Israel |
| Human single-chromosome paint probes,  Chromosome 18 Green (FITC) | Custom-synthesised | Applied Spectral Imaging, Israel |
| Human single-chromosome paint probes,  Chromosome 22 Green (FITC) | Custom-synthesised | Applied Spectral Imaging, Israel |
| Human Pan-Centromeric probes | Custom-synthesised | Chrombios GmbH, Germany |
| Pan-Telomeric probe  G-rich probe (repeats of TTAGGG) | F1006 | HLB Panagene Co., Ltd, Republic of Korea |
| Human 18S ribosomal RNA probe 5′ 6-FAM  (5′-UUGAGACAAGCAUAUGCUACCUGGC-3′) | Custom-synthesised | Integrated DNA Technologies, USA |
| Human LINE1 FISH probe (40 mer) 5′ 6-FAM  (5′-CACGCCATTGCACTCCAGCC  TGAGCTATGGGAGTAAAACC-3′) | Custom-synthesised | Eurofins Genomics, India |
| Human Alu Elements probe (30 mer) 5’ 6-FAM  (5’-CAAACACACACCACCACACCCAACTAATTT-3’) | Custom- synthesised | Eurofins Genomics, India |

**Non-coding RNA probe:**

| **Probes** | **Catalogue no.** | **Company** |
| --- | --- | --- |
| Stellaris® FISH Probes, Human LSINCT5 with FAM | VSMF-20343-5 | LGC Biosearch Technologies, UK |
| Stellaris® FISH Probes, Human LSINCT5 with Quasar 570 | VSMF-20339-5 | LGC Biosearch Technologies, UK |
